# A mutation in the transmembrane domain of *Adenylate cyclase 3* impairs enzymatic function to cause sex-specific depression- and anxiety-like behaviors and food seeking in a rat model

**DOI:** 10.1101/2025.03.28.645767

**Authors:** Mackenzie K. Fitzpatrick, Christina Dyson, Angela Beeson, Leighelle Adrian, Glen Marrs, Michael Grzybowski, Jason Klotz, Aron M. Geurts, Rong Chen, Jeffrey L. Weiner, Leah C. Solberg Woods

## Abstract

We have previously demonstrated that a transmembrane domain mutation in *Adenylate cyclase 3* (*Adcy3*) causes increased adiposity and negative emotion-like behaviors in a rat model. We set out to replicate and expand upon our previous study by conducting comprehensive behavioral testing, and we also investigated the molecular changes that result from this mutation.

Rats with a mutation in the second transmembrane helix of ADCY3 (Adcy3^mut/mut^) and wild-type rats were fed a high-fat diet for 12 weeks. We measured body weight, body composition, and depression-like and anxiety-like behaviors using the following tests: sucrose splash test, sucrose preference test, forced swim test, open field test, elevated plus maze, successive alleys test, and novelty-suppressed feeding. We also measured serum leptin levels, hypothalamic cyclic AMP (cAMP) production, and membrane fraction ADCY3 content.

Adcy3^mut/mut^ male and female rats had increased adiposity. Adcy3^mut/mut^ males showed increased despair- and anxiety-like behaviors, food seeking, and higher leptin levels relative to wild-type males. Adcy3^mut/mut^ females showed only mildly increased anxiety-like behaviors relative to wild-type females. Adcy3^mut/mut^ rats of both sexes had decreased cAMP production in the hypothalamus, with no changes in ADCY3 content in the membrane fraction.

We conclude that the transmembrane domain of ADCY3 plays a critical role regulating adiposity and behavior, as well as cAMP production. There were key differences between males and females for the observed phenotypes. This study supports the idea that *Adcy3* contributes to emotion-like behaviors and potentially mental health disorders, and that the transmembrane domain of ADCY3 is important for protein function.

## 2 Introduction

Major depressive disorder (MDD) prevalence has been rapidly increasing over the last few decades, with ∼21 million adults in the US experiencing at least one major depressive episode in 2021.^1, 2^ The prevalence of anxiety in the US is also increasing, particularly in adults under age 50.^3^ Both environmental and biological factors contribute to a person’s risk of developing mental health disorders.^4, 5^ Genetic factors have also been shown to play a role in both MDD and anxiety, although identification and validation of specific candidate genes has been limited.^6, 7^ Furthermore, MDD in particular has been shown to be bi-directionally associated with multiple physical comorbidities, including heart disease, osteoarthritis, metabolic syndrome, and obesity.^8–13^

Obesity rates are steadily increasing as well, and it has been predicted that 51% of the global population will be overweight or obese by 2035.^14^ Like MDD and anxiety, genetic risk factors have been identified in association with obesity, with more than 500 candidate genes identified thus far.^15, 16^ Some of these gene have also been identified as candidate genes for MDD,^17, 18^ further demonstrating the important association between these two diseases. Additionally, shared underlying biological pathways have been identified for all three diseases, and impaired cyclic AMP (cAMP) signaling in particular has been linked to MDD, anxiety, and obesity.^19–22^ cAMP is a secondary messenger that plays a crucial role in hormone secretion, gene expression, and neuronal signaling,^23^ and both antidepressants and obesity treatment medications have been shown to increase cAMP concentrations.^21, 24^ Despite the fact that MDD and obesity have been shown to be bi-directionally associated, and that anxiety may also share causal mechanisms with MDD and obesity, very few studies have investigated biological and genetic mechanisms that may contribute to these associations.

The gene *Adenylate cyclase 3 (Adcy3)* has been identified as a candidate gene for both MDD^20, 25, 26^ and obesity.^27, 28^ Upon activation, the ADCY3 protein catalyzes the synthesis of cAMP. *Adcy3* is highly expressed in the brain, particularly in the hypothalamus, hippocampus, and olfactory regions.^29^ Multiple *Adcy3* variants have been associated with obesity in humans,^28^ and ADCY3 has also been proposed as a blood biomarker for MDD.^20, 25, 26^ These associations are supported by *Adcy3* mouse knockout (KO) models, where KOs show increased body weight, fat mass, food intake, and depression-like and anxiety-like behaviors relative to wild-type (WT) mice.^30–35^

Our lab previously identified^36^ and validated^37^ a protein-coding variant (Adcy3^mut/mut^) in the transmembrane (TM) domain of *Adcy3* that causes obesity in a rat model. The TM domain of ADCY3 has been shown to play an important role in both membrane localization^38^ and catalytic activity.^39^ Similar to the mouse knock-out models,^30–33^ we found that Adcy3^mut/mut^ rats showed both increased adiposity and increased negative emotion-like behaviors compared to WT rats, demonstrating the importance of the TM domain for protein function and supporting the idea that *Adcy3* may serve as a genetic link between obesity and MDD and/or anxiety.^37^ However, we also unexpectedly observed sex differences in these phenotypes. The underlying cause of adiposity in Adcy3^mut/mut^ differed by sex, where Adcy3^mut/mut^ males, but not females, consumed more food than their WT counterparts. In addition, Adcy3^mut/mut^ males showed increased despair-like behavior in the forced swim test (FST), decreased memory in the novel object recognition test (NOR) and altered behavior in the open field test (OFT), while Adcy3^mut/mut^ female rats only exhibited increased anxiety-like behavior in the OFT with no differences in the FST or NOR.^37^ We found that decreases in downstream signaling proteins, namely PKA, AMPK, and CREB, may be causing the adiposity and behavior phenotypes, and we observed sex differences in PKA and CREB signaling as well.^37^

In the present study, we set out to better understand the nuances of the altered behavior observed in Adcy3^mut/mut^ male and female rats. We performed comprehensive behavioral analyses of Adcy3^mut/mut^ to replicate and expand upon the testing performed in the previous study. We also investigated the molecular changes caused by Adcy3^mut/mut^ to better understand how this mutation impacts ADCY3 protein function.

## 3 Materials and Methods

### 3.1 Animals and Diet

Adcy3^mut/mut^ rats were generated, genotyped, and housed as previously described.^37^ Briefly, we used CRISPR-SpCas9 to generate rats that have a protein coding mutation (F122delV123L) in the second ADCY3 TM helix on the Wistar Kyoto (WKY) background. We used the WKY strain for this model because our initial goal was to assess the effects of a related *Adcy3* variant (L121P) that we previously identified in WKY rats.^36^ CRISPR errors instead generated Adcy3^mut/mut^, which we investigated instead as it is still a model of an adiposity-associated TM-domain *Adcy3* mutation. Heterozygous breeding pairs were used to establish a breeding colony at the Wake Forest University School of Medicine (WFUSOM). 12 rats per genotype-sex group were collected for behavioral experiments. Additional cohorts collected for the sucrose preference test resulted in N=22-24 rats per group for WT and Adcy3^mut/mut^ male rats in that test only. A separate group of rats (10-12 per group) was used to collect serum for measuring leptin content, and hypothalamus samples (3 per group) were collected from this same group for high-fat diet (HFD) membrane fractionation experiments. A third group of rats (6-8 per group) was used to collect chow-diet hypothalamus samples for cAMP activity assays and membrane fractionation. A fourth group of rats (2-3 per group, males only) was used to collect samples for immunofluorescence.

We used Adcy3^+/-^ rats for visual comparison in immunofluorescence experiments; these rats were also generated, genotyped, and housed as previously described.^37^ In summary, we generated *Adcy3* KO rats that have single base pair deletion that causes a frameshift and full KO of *Adcy3* on the WKY background, and we used Adcy3^+/-^ rats as experimental animals from this strain due to homozygous lethality in *Adcy3* KO rats.^37^ Previous work by our group has shown that both Adcy3^+/-^ and Adcy3^mut/mut^ male and female rats exhibit increased adiposity when on a HFD relative to WT rats.^37^ 2-3 rats per group (males only) were used to collect samples for immunofluorescence.

All rats were housed at standard temperature and humidity conditions on a 12-hour light/dark cycle with *ad libitum* access to food and water. All testing and data collection was performed during the light phase. Food for all rats up to 5 weeks of age consisted of standard rodent chow (Lab Diet, Prolab RMH 3000, #5P00, St. Louis, MO). In keeping with the previous study of Adcy3^mut/mut^,^37^ and because prior work had shown that HFD is required for the adiposity phenotype in *Adcy3* KO mice,^30^ all rats used for behavioral experiments, serum leptin measurement, some membrane fractionation experiments (**Figure S3**), and immunofluorescence experiments began a high-fat diet (HFD) (60% kcal fat, 20% kcal protein, 20% kcal carbohydrates, ResearchDiet D12492, New Brunswick, NJ) at 5 weeks of age. Rats used for cAMP activity assays and other membrane localization assays (**Figure 9**) were maintained on standard chow. All animal experiments were performed using protocols approved by the Institutional Animal Care and Use Committee at WFUSOM.

### 3.2 Study Design

Rats were weaned at 3 weeks of age and pair housed. Beginning at HFD start, body weight was measured weekly. Behavioral phenotyping began at 6 weeks on diet (WOD) (**Figure 1A**). Rats received an EchoMRI and 3 behavior tests (splash test, sucrose preference test (SPT), and forced swim test (FST)) in Building A before being transferred to Building B for 4 additional tests (open field test (OFT), elevated plus maze (EPM), successive alleys test (SAT), and novelty-suppressed feeding (NSF)) (**Figure 1A**). Rats were given 10 days to acclimate to Building B before any behavior tests were conducted. Caging, lighting conditions, and diet were consistent across buildings.

**Figure 1.**
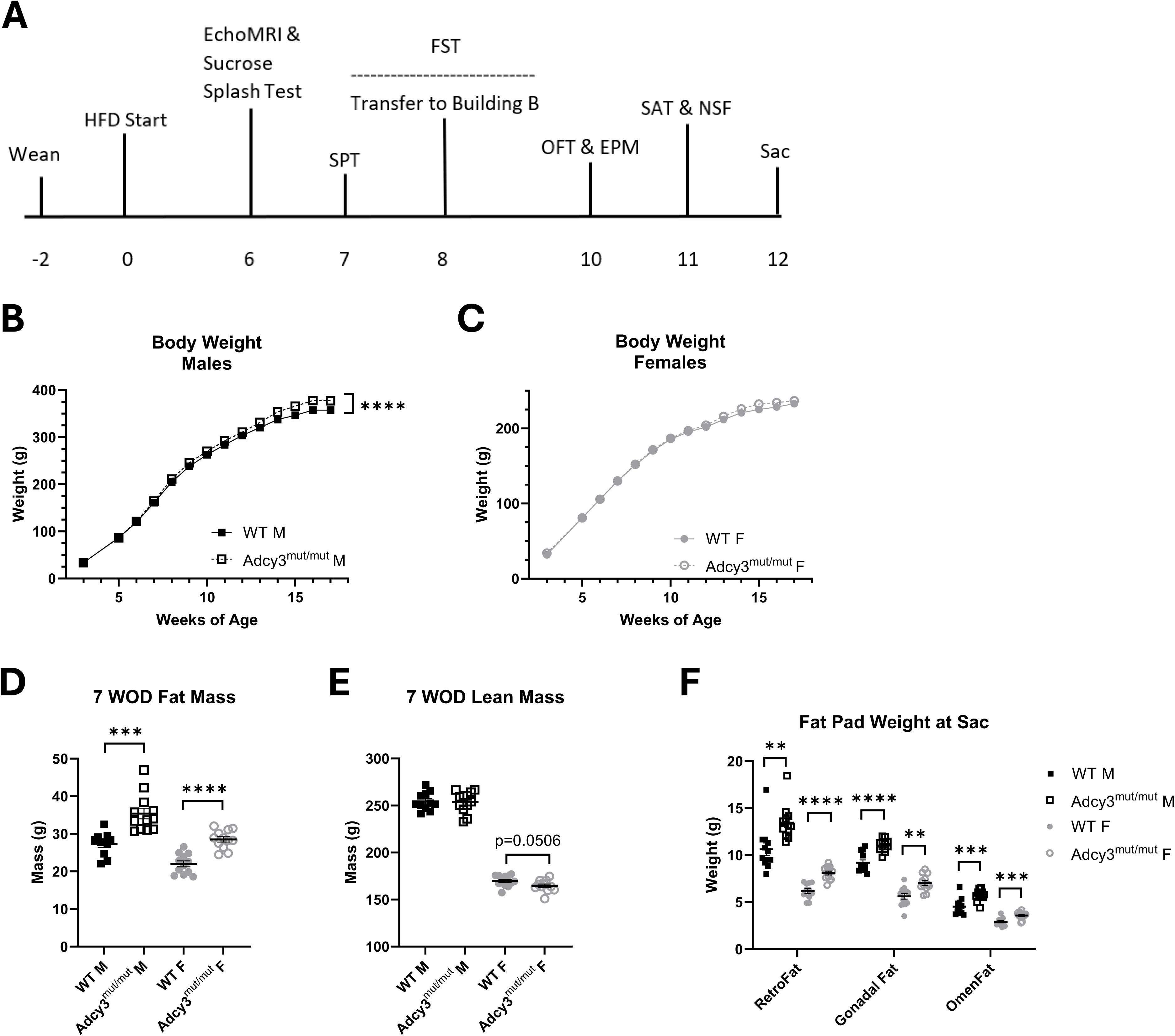
Study design, body weight, and fat and lean mass in Adcy3^mut/mut^. **(A)** Timeline of the study in weeks on diet (WOD). Rats were given an EchoMRI and 7 behavioral tests across 2 buildings before necropsy (sac) at 12 WOD. **(B)** Adcy3^mut/mut^ males (M) gain more weight than wild-type (WT) M over the course of the study. **(C)** There were no significant differences in body weight between Adcy3^mut/mut^ and WT females (F). **(D)** After 7 WOD, both Adcy3^mut/mut^ M and F have more fat mass than WT rats. **(E)** Adcy3^mut/mut^ F tend to have less lean mass than WT F. **(F)** Adcy3^mut/mut^ M and F have larger retroperitoneal (RetroFat), gonadal, and omental (OmenFat) fat pads at sac than WT. SPT: sucrose preference test, FST: forced swim test, OFT: open field test, EPM: elevated plus maze, SAT: successive alleys test, NSF: novelty-suppressed feeding test. Mean ± SEM. T-test or rmANOVA, **p<0.01, ***p<0.001, ****p<0.0001

### 3.3 EchoMRI

Rats were weighed, then fat mass and lean mass were measured in triplicate (EchoMRI LLC, Houston, TX).

### 3.4 Behavioral Phenotyping

Before each behavior test, rats were given at least 30 minutes to acclimate to the testing room. All behavior was either automatically scored in Ethovision 15 (Noldus, Leesburg, VA) (OFT, EPM, SAT) or video-recorded and scored by an experimenter blind to the rat’s group (splash test, FST, NSF). SPT intake was measured manually as described below. Any shared items or arenas were cleaned between rats with Quatricide PV-15 (Pharmacal, Jackson, WI).

#### 3.4.1 Sucrose Splash Test

Rats received the sucrose splash test after 6 WOD to assess motivational and self-care behaviors, which are components of depression-like behavior.^40, 41^ Each rat was tested individually in a clean cage that was identical to the home cage, but without food and water. After acclimating to the clean cage for 10 minutes, each rat was sprayed on their dorsal coat with a 10% sucrose solution. Over 5 minutes, latency to groom, total grooming time, and grooming time within each 1-minute bin were scored.

#### 3.4.2 Sucrose Preference Test (SPT)

Rats received the SPT after 7 WOD to assess anhedonia, which is also a component of depression-like behavior.^40, 41^ Rats were individually housed with 2 water bottles in each cage. After 24 hours of acclimation, both bottles were filled with a 1% sucrose solution, and rats were again given 24 hours to acclimate. Rats were then given one bottle of 1% sucrose and one bottle of water. Sucrose intake and water intake over 24 hours were measured, with the positions of the bottles switched halfway through the measurement period. The initial position of the sucrose bottle was counterbalanced across genotypes. If either bottle leaked noticeably onto the cage floor, the rat was excluded from analysis (N=1 WT male and N=2 WT females).

#### 3.4.3 Forced Swim Test (FST)

Rats received the FST after 8 WOD as previously described^37^ to assess passive coping/depression-like behavior.^42^ Briefly, rats were placed in a cylindrical tank (30 cm diameter x 50 cm tall) of 25 water and behavior was scored at 5-second intervals as “mobile” (swimming, climbing, or diving) or “immobile”.

#### 3.4.4 Open Field Test (OFT)

Rats received the OFT after 10 WOD to assess anxiety-like behavior.^41^ Each rat was placed in a standard OFT chamber (42 cm x 42 cm x 30 cm, Omnitech, Sioux Falls, SD) for 30 minutes. The chamber was illuminated by a 7.5-watt light bulb mounted 48 cm over the arena. Multiple behaviors were scored over 30 minutes, with scoring binned in 5-minute intervals. Behaviors scored included total distance traveled, center time, grooming activity, and vertical rearing time. Grooming activity was measured by beam breaks, where the Ethovision 15 software considers multiple repeated breaks of the same beam to be repetitive behavior associated with grooming.^43^

#### 3.4.5 Elevated Plus Maze (EPM)

Rats received the EPM after 10 WOD as an additional test of anxiety-like behavior.^40, 41^ Each rat was placed in a standard EPM chamber (Med Associates, Fairfax, VT) consisting of two open arms and two closed arms (each arm is 10.2 cm x 50.8 cm). The chamber was illuminated at approximately 0 lux in each closed arm and 90 – 120 lux for each open arm. Each rat was placed in the junction of the chamber facing the open arm, and entries into and time spent in each arm were recorded over 5 minutes.

#### 3.4.6 Successive Alleys Test (SAT)

Rats received the SAT after 11 WOD as an additional test of anxiety-like behavior.^44^ The SAT apparatus has four segments (each 45 cm in length) that are progressively brighter and more exposed: enclosed (9.0 cm wide with 29.0 cm walls), dark gray (9.0 cm wide with 2.5 cm walls), light gray (6.7 cm wide with 0.5 cm walls), and white (3.5 cm wide with 0.3 cm walls). The apparatus is illuminated by a 60-watt light bulb mounted 120 cm above the apparatus. Each rat was placed in the enclosed segment, and behavior was recorded over 5 minutes. Behaviors scored included latency to enter, duration in, and frequency in each segment as well as average velocity and total distance traveled.

#### 3.4.7 Novelty-Suppressed Feeding (NSF)

Rats received the NSF after 11 WOD as an additional test of anxiety-like and/or depressive-like behaviors.^40, 41^ Female rats were fasted for 24 hours prior to testing while male rats were fasted for 48 hours. A HFD food pellet was placed in the center of an NSF chamber (67.6 cm x 86.4 cm). The chamber was illuminated by four bright light bulbs, one at each corner of the arena, resulting in approximately 120 – 165 lux in the center of the area and 40 – 50 lux in the margins. A small amount of peanut butter was added to the food pellet for males. The increased fasting time and addition of the peanut butter was required to entice the males to participate in the test. On the day of the test, each rat was placed in the corner of the arena and allowed to freely move and eat for 10 minutes. Rats that did not eat at all during the test were excluded from analysis (N=5 WT males, N=4 Adcy3^mut/mut^ males, N=1 WT female, N=2 Adcy3^mut/mut^ females). Eating frequency, duration, and latency as well as food approach frequency and latency were scored. “Frequency” was defined as the total number of eating episodes or approaches over 10 minutes.

### 3.5 Tissue Harvest

Rats from behavioral experiments were euthanized via live decapitation after 12 WOD following a 4-hour fast. Rats collected for other purposes had no fast before euthanasia. The following tissues were weighed and snap-frozen in liquid nitrogen: brain, pituitary, liver, retroperitoneal fat (RetroFat), adrenal glands, gonadal fat, gonads, spleen, brown adipose (BAT), subcutaneous adipose (SubQ), and tail. Kidneys, omental fat (OmenFat), pancreas, and heart were weighed only.

### 3.6 Serum Leptin

Serum was collected from a separate group of rats via tail snip after a 16-hour overnight fast. Serum leptin was measured with a rat leptin enzyme-linked immunosorbent assay (ELISA) kit (Crystal Chem, #90040, Elk Grove Village, IL). Leptin concentrations were interpolated from optical density values using 4-parameter logistic regression in GraphPad Prism (10.2.2, Boston, MA).

### 3.7 cAMP Activity Assay

cAMP production in a third group of rats was measured as previously described.^30^ Briefly, membrane protein was collected from hypothalamus samples using ultracentrifugation in a buffer containing HEPES and sucrose.^30^ Protein concentrations were measured by bicinchoninic acid assay (Thermo Scientific, #23225, Waltham, MA) and standardized to 1.25μg/μl. 200μl of standardized protein was combined with 100μl of “reaction mixture”, a Tris-acetate buffer containing multiple components including ATP, MgCl_2_, 3-isobutyl-1-methylxanthine, and either forskolin or no forskolin.^30^ The mixture was incubated in a water bath at 37°C for 30 minutes. Reaction was terminated at 95°C for 5 minutes and mixtures were centrifuged at 10,000xg for 5 minutes, then supernatants containing cAMP were collected. cAMP concentrations were measured using a cAMP ELISA kit (Enzo Life Sciences, #ADI-900-067A, Farmingdale, NY). cAMP concentrations were interpolated from optical density values using 4-parameter logistic regression in GraphPad Prism.

### 3.8 Membrane Fractionation and Western Blot

Protein was extracted from hypothalamus samples in a 50mM Tris buffer (150mM NaCl, 10% sucrose, 3μl protease inhibitor cocktail (Sigma Aldrich, #P8340, St. Louis, MO)) using a Bead Beater Tissue Homogenizer at 2200rpm with 1.0mm zirconia/silica beads (BioSpec, #11079110Z, Bartlesville, OK). Homogenates were centrifuged at 20,000xg at 4°C for 1 hour. Supernatant containing cytosolic proteins was collected. Pellet was resuspended in a 30mM Tris buffer (100mM NaCl, 1% sodium cholate, 3μl protease inhibitor cocktail) with shaking for 1 hour at 4°C. Samples were then centrifuged at 100,000xg at 4°C for 1 hour, and supernatant containing membrane proteins was collected. Western blot was then performed as previously described.^37^ Primary antibodies to ADCY3 (EnCor, RPCA-ACIII, 1:2000, Gainesville, FL) and α-tubulin (CST, #2144, 1:2000, Danvers, MA) were used.

### 3.9 Immunofluorescence

A separate group of rats (Adcy3^mut/mut^ and Adcy3^+/-^) were transcardially perfused with phosphate-buffered saline then with 4% paraformaldehyde (PFA). Brains were extracted and fixed in 4% PFA overnight followed by cryoprotection in 30% sucrose. Brains were then frozen in Optimal Cutting Temperature compound (Fisher Healthcare, #4585, Waltham, MA), sliced in 30μm coronal sections through the hypothalamus, and mounted on glass slides. Slides were processed through antigen retrieval in a citrate-based solution (Vector Labs, #H-3300-250) at 80°C for 30 minutes before blocking for 2 hours in 10% Normal Goat Serum (NGS) (Thermo Fisher, #PCN5000, Waltham, MA) + 0.3% Triton X-100 (Sigma Aldrich, #X100-100ML, St. Louis, MO). Slides were incubated overnight at 4°C with primary antibody to ADCY3 (EnCor, RPCA-ACIII, 1:5000, Gainesville, FL) diluted in 10% NGS. After washing, slides were incubated with secondary antibody (Thermo Fisher, #A21207, Waltham, MA) diluted in 10% NGS for 90 minutes at room temperature before washing again and mounting with Prolong Gold with 4’,6-diamidino-2-phenylindole (DAPI) (Thermo Fisher, #P36941, Waltham, MA).

Confocal images were captured with the 40X Oil (1.3 NA) objective on a Zeiss LSM 710 microscope (Oberkochen, Germany) using 594nm and 405nm excitation laser lines. Identical sample excitation and acquisition settings were used for all images. Maximum intensity projections were generated from z-stacks with 10 slices spanning 10.8μm. Image processing was done in ImageJ (version 1.54g, NIH, Bethesda, MA) where brightening and contrast adjustments were applied identically to maximize image visualization.

### 3.10 Statistical Analysis

Data were analyzed in RStudio (2023.09.1, Boston, MA) and GraphPad Prism. Outliers were identified by Grubb’s Test or by the ROUT method in Prism with Q=1% (if multiple outliers suspected). In the sucrose splash test, rats that were outliers for “latency to groom” were removed from all variables. In the OFT, a rat was only removed from a variable if they were an outlier in at least 3 time bins for that variable. If removing outliers would result in removing all or most non-zero values, outliers were left in.

Data were assessed for normality with the Shapiro-Wilk test and, if not normal, Mann-Whitney U test was used in place of a Student’s t-test for analysis. Sphericity was assessed with Mauchly’s Test for repeated measures analysis of variance (rmANOVA), and Greenhouse-Geisser correction was applied as necessary.

Data were first analyzed using a two-way ANOVA or rmANOVA with genotype and sex as main factors. rmANOVA analyses were used for body weight, splash test, and OFT data. Due to the presence of significant sex effects, all data were also stratified by sex and analyzed separately using an unpaired Student’s t-test or a one-way rmANOVA as appropriate for the structure of the data collected. We only report stratified analyses for adiposity measures due to large sex differences in these phenotypes.

## 4 Results

### 4.1 Adcy3^mut/mut^ rats weigh more than WT rats and have more fat mass

Adcy3^mut/mut^ males weighed significantly more than WT males (F_13,273_=8.31, p<0.0001) (**Figure 1B**). Adcy3^mut/mut^ females tended to weigh more than WT females, especially near the end of the study, although the difference was not significant (**Figure 1C**). After 7 WOD, both Adcy3^mut/mut^ males (t_21_=4.64, p=0.0001) and females (t_22_=6.21, p<0.0001) had significantly more fat mass than WT rats (**Figure 1D**). There were no significant differences in lean mass in Adcy3^mut/mut^ males (**Figure 1E**). However, Adcy3^mut/mut^ females showed a slight decrease in lean mass that was nearly significant (t_22_=2.07, p=0.0506; **Figure 1E**), potentially explaining why there was not a significant body weight difference between WT and Adcy3^mut/mut^ females despite the large difference in fat mass. Adcy3^mut/mut^ males and females both had significantly more RetroFat (t_22_=3.32, p=0.0031; t_21_=5.89, p<0.0001), gonadal fat (t_22_=5.22, p<0.0001; t_22_=3.59, p=0.0016), and OmenFat (t_22_=3.93, p=0.0007; t_22_=4.07, p=0.0005) than WT rats at necropsy (**Figure 1F**). There were no significant differences in the necropsy weights of any other organs (**Figure S1**).

### 4.2 Adcy3^mut/mut^ males groom more than WT males in the sucrose splash test with no genotypic differences in females

In the sucrose splash test, there were no significant genotypic differences in latency to groom or total grooming time over the full five minutes of the test, although there was a significant sex effect: males groomed for significantly more time than females (F_1,42_=4.25, p=0.0455) (**Figure 2A-B**). When stratified by sex and binned to generate grooming time curves, Adcy3^mut/mut^ males showed a significantly different pattern of grooming compared to WT males (F_4,84_=2.79, p=0.0312), where Adcy3^mut/mut^ males groom more than WT males during the first 2 minutes of the test (**Figure 2C**). There were no significant differences between the grooming time curves of Adcy3^mut/mut^ and WT females (**Figure 2D**).

**Figure 2.**
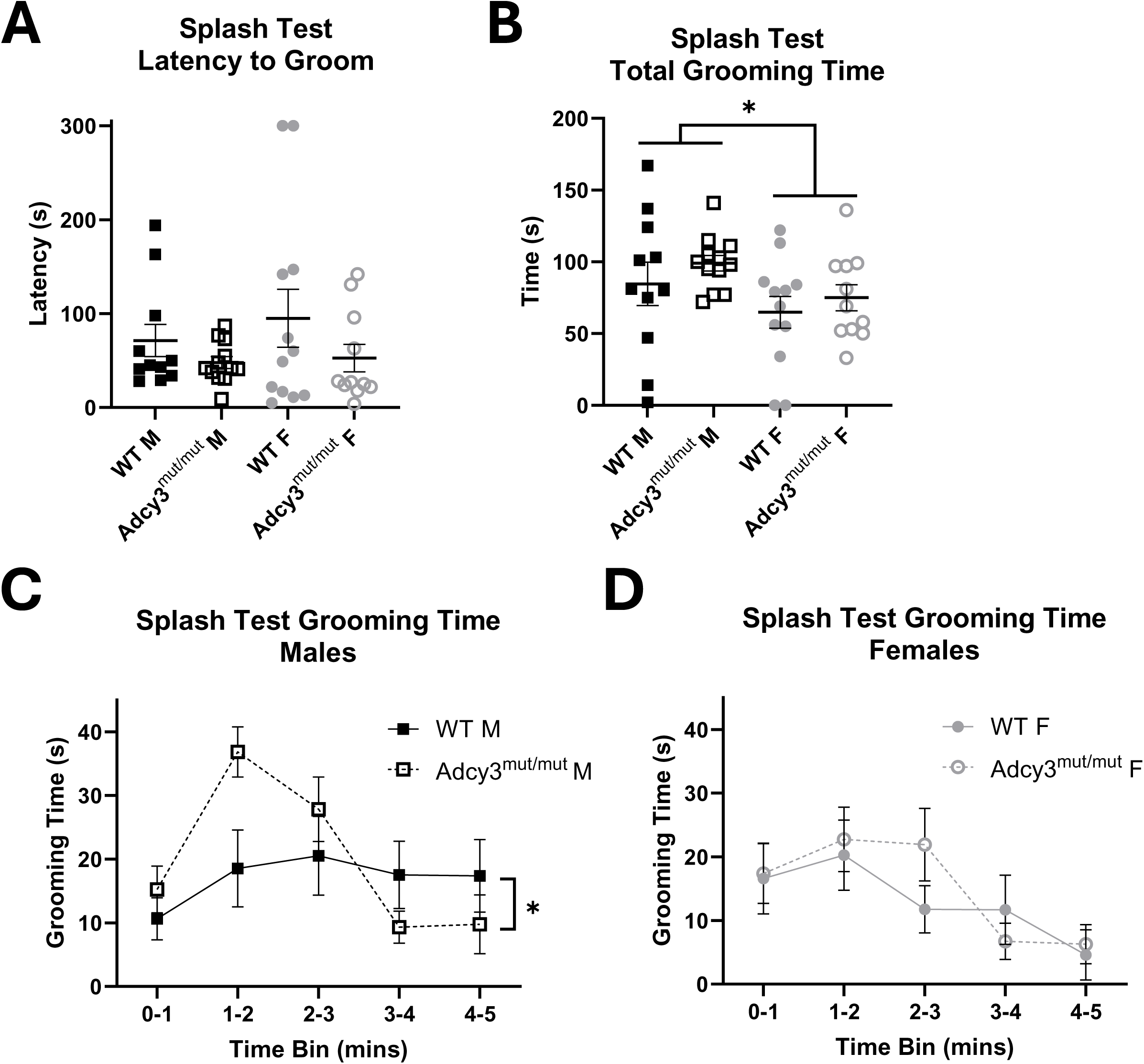
Adcy3^mut/mut^ behavior in the sucrose splash test. **(A)** No differences between Adcy3^mut/mut^ and wild-type (WT) rats in splash test latency to groom. **(B)** No differences between Adcy3^mut/mut^ and WT rats in splash test total grooming time over the full five minutes, although males (M) did groom significantly more than females (F). **(C)** Adcy3^mut/mut^ M show a significantly different pattern of grooming in the splash test relative to WT M, grooming more than WT M in the first 2 minutes of the test. **(D)** No differences between Adcy3^mut/mut^ F and WT F in splash test grooming. Mean ± SEM. Two-way ANOVA or rmANOVA, *p<0.05

### 4.3 WT and Adcy3^mut/mut^ rats prefer sucrose over water

In the sucrose preference test, all four groups significantly preferred the sucrose solution over water (t_22_=21.60, p<0.0001; t_21_=17.60, p<0.0001; t_9_=12.89, p<0.0001; t_10_=20.02, p<0.0001) at approximately an 80% sucrose preference rate (**Figure 3**). However, there were no significant sex, genotype, or interaction effects for sucrose preference rate.

**Figure 3.**
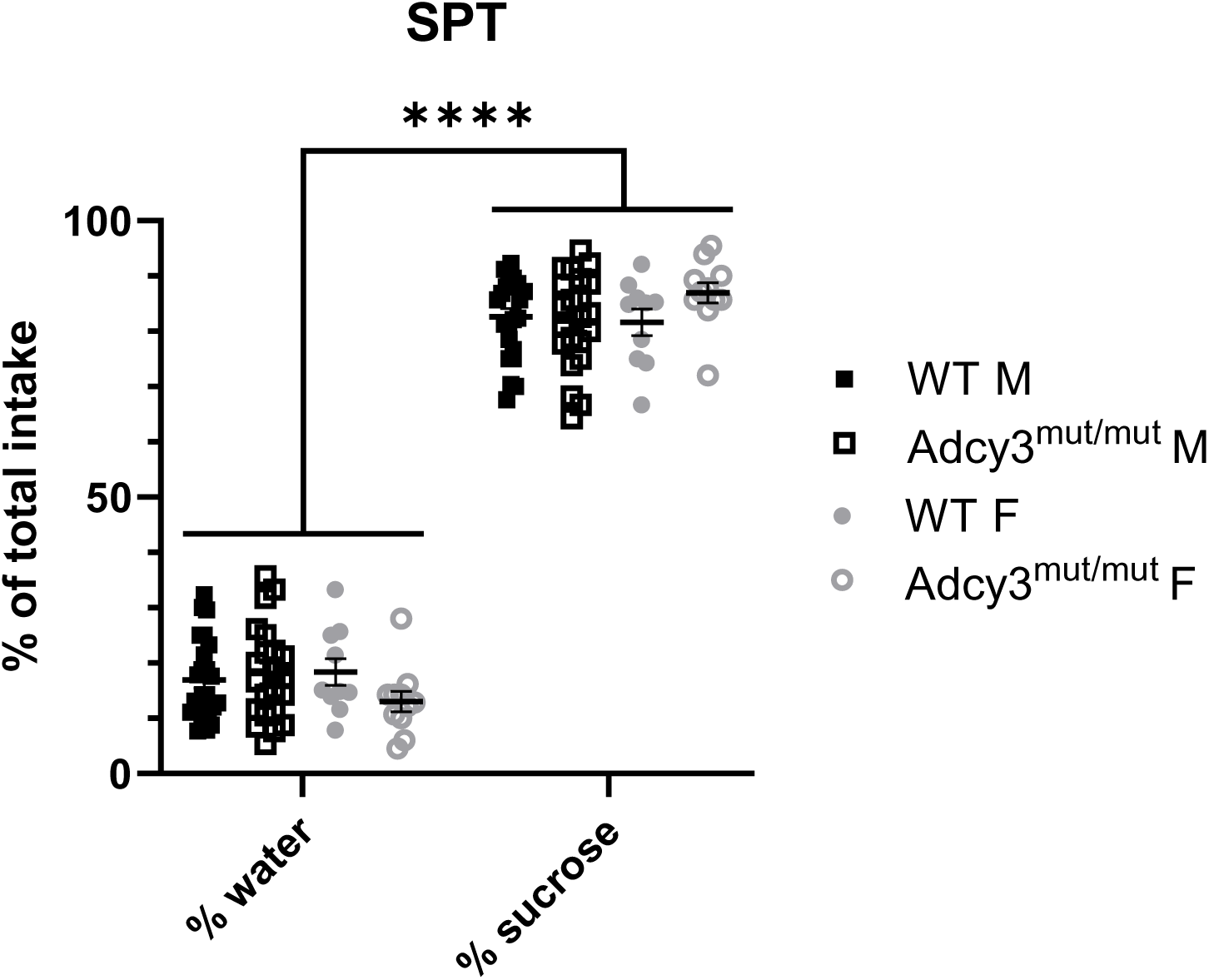
Adcy3^mut/mut^ behavior in the sucrose preference test (SPT). All four groups significanlty preferred sucrose over water, but there were no significant differences between Adcy3^mut/mut^ and WT rats in sucrose preference rate. Mean ± SEM. Two-way ANOVA, ****p<0.0001

### 4.4 Adcy3^mut/mut^ males are significantly more immobile in the FST with no differences in females

In the FST, there were significant sex (F_1,43_=8.09, p=0.0068) and genotype by sex interaction (F_1,43_=5.70, p=0.0214) effects on both mobile and immobile counts (**Figure 4**). When separated by sex, Adcy3^mut/mut^ males spent significantly more time immobile and significantly less time mobile in the FST than WT males (t_21_=2.77, p=0.0115), indicating increased passive coping and/or depression-like behavior (**Figure 4**). There were no significant differences in mobile or immobile counts in Adcy3^mut/mut^ females relative to WT females (**Figure 4**).

**Figure 4.**
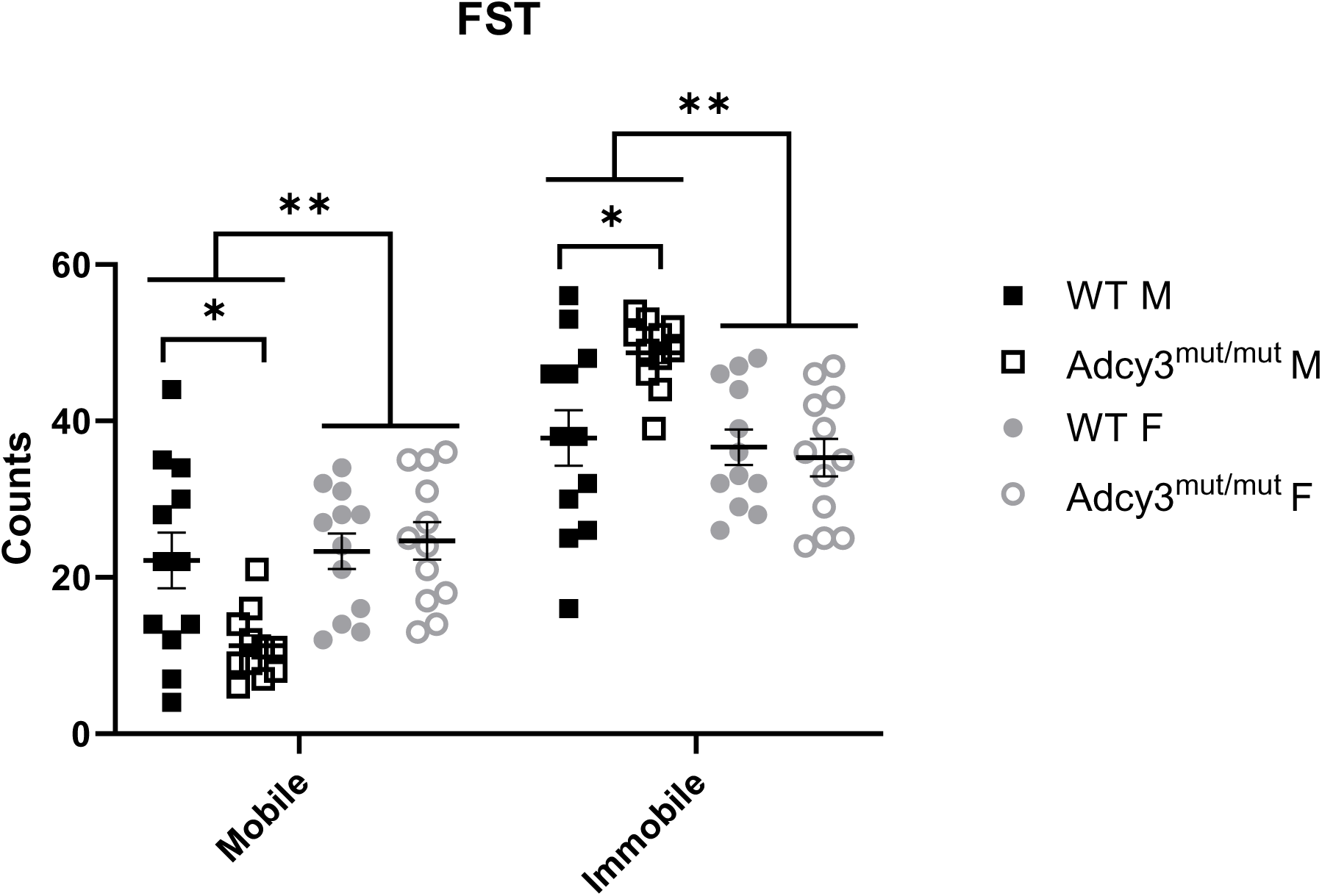
Adcy3^mut/mut^ behavior in the forced swim test (FST). Adcy3^mut/mut^ males (M) were less mobile and more immobile in the FST than wild-type (WT) M. There were no differences between Adcy3^mut/mut^ and WT females (F) in the FST, although F were significantly more mobile than M. Mean ± SEM. Two-way ANOVA or T-test, *p<0.05, **p<0.01

### 4.5 Adcy3^mut/mut^ males and females show increased anxiety-like behaviors in the OFT

In the OFT, there was a significant effect of sex on total distance traveled (F_1,42_=74.31, p<0.0001), where females traveled further than males during the test (**Figure 5A**). Due to this large difference in total motion, we stratified the data for all other OFT variables by sex and conducted analyses separately for males and females.

**Figure 5.**
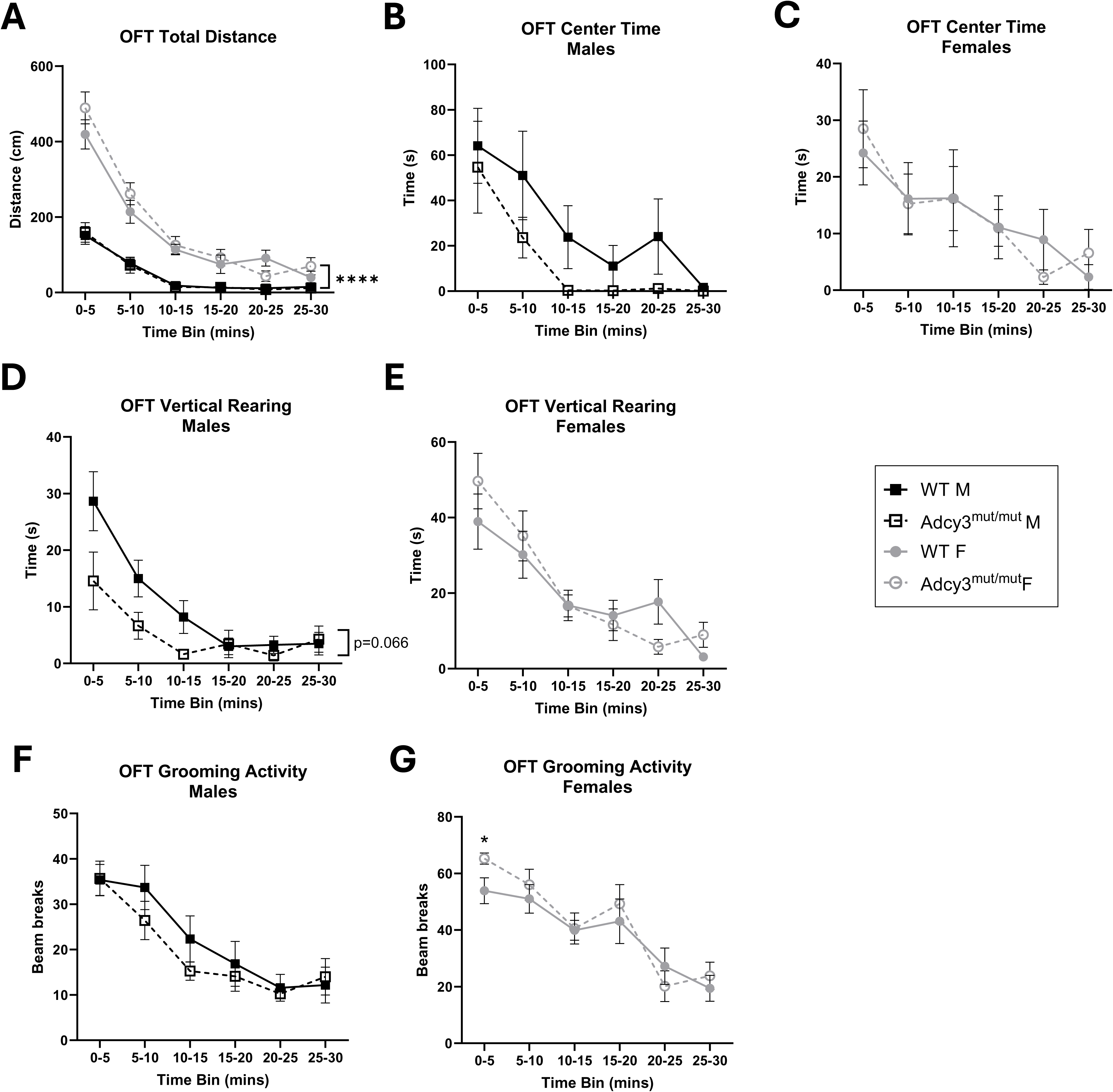
Adcy3^mut/mut^ behavior in the open field test (OFT). **(A)** Females (F) have increased OFT total distance relative to males (M), although there are no differences between Adcy3^mut/mut^ and wild-type (WT) rats in OFT total distance. **(B)** Adcy3^mut/mut^ M appear to have decreased OFT center time, although this does not reach significance. **(C)** No differences between Adcy3^mut/mut^ and WT F in OFT center time. **(D)** Adcy3^mut/mut^ M spend less time vertically rearing in the OFT than WT M. **(E)** No differences between Adcy3^mut/mut^ and WT F in OFT vertical rearing. **(F)** No differences between Adcy3^mut/mut^ and WT M in OFT grooming activity. **(G)** Adcy3^mut/mut^ F groom more than WT F in the first 5 minutes of the OFT. Mean ± SEM. T-test, two-way ANOVA, or rmANOVA, *p<0.05, ****p<0.0001

There were no differences by genotype in total distance traveled in either sex (**Figure 5A**). Adcy3^mut/mut^ males appear to have decreased center time relative to WT males, although this does not reach significance (**Figure 5B**). There were no significant differences in center time by genotype in females (**Figure 5C**). Adcy3^mut/mut^ males also tended to spend less time rearing than WT males (F_5,100_=2.76, p=0.0661), especially in the first 15 minutes of the test (**Figure 5D**). There were no differences in rearing time in Adcy3^mut/mut^ females compared to WT females (**Figure 5E**). Adcy3^mut/mut^ females did show significantly more grooming activity than WT females in the first 5 minutes of the test (t_22_=2.27, p=0.0334) (**Figure 5G**). There were no significant differences in grooming activity in Adcy3^mut/mut^ males compared to WT males (**Figure 5F**). Decreased center time, decreased rearing, and increased grooming are all consistent with an increase in anxiety-like behaviors, although the magnitude of this increase is modest in both sexes.

### 4.6 Adcy3^mut/mut^ rats do not show any differences in behavior in the EPM or SAT

In the EPM, there were no genotype or interaction effects for time spent in each arm type in the two-way ANOVA or in the stratified analysis (**Figure 6A**). There were also no genotype or interaction effects for closed arm or open arm entries (**Figure 6B-C**). There was a significant sex effect for closed arm time (F_1,43_=9.94, p=0.0029) and closed arm entries (F_1,43_=7.92, p=0.0074) in the two-way ANOVA, where female rats entered the closed arms more often and spent more time in them than male rats (**Figure 6A-B**). However, rats in all groups spent the most time in the junction area of the arena during the test, and the total number of entries into either arm type was relatively low for both sexes (**Figure 6A-C**).

**Figure 6.**
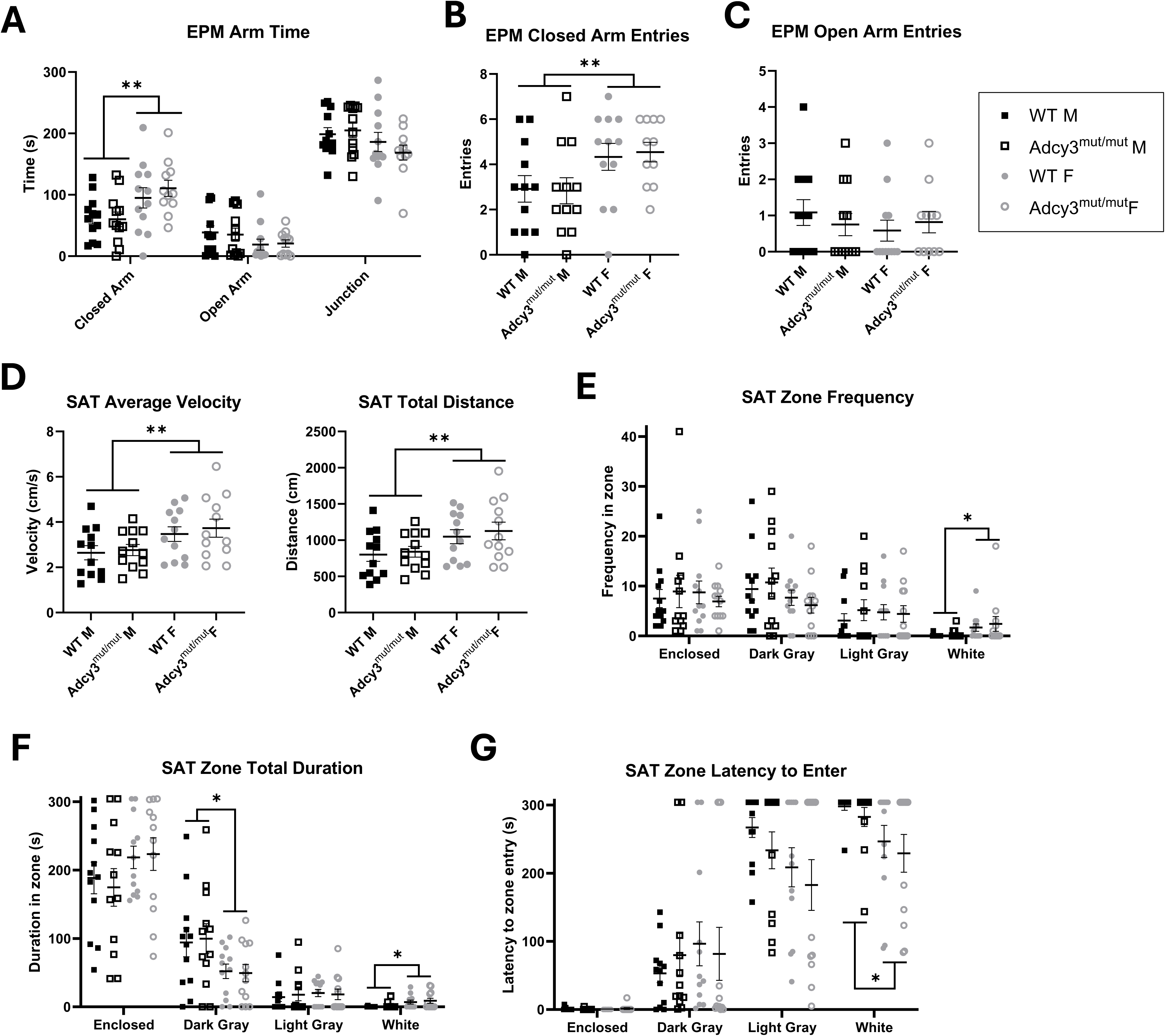
Adcy3^mut/mut^ behavior in the elevated plus maze (EPM) and successive alleys test (SAT). **(A-G)** No differences between Adcy3^mut/mut^ and wild-type (WT) rats in any variable measured in the EPM or SAT, although there are sex differences in several variables. Both WT and Adcy3^mut/mut^ rats spend most of their time in the EPM junction and the enclosed arm of the SAT, suggesting a floor effect is present. Mean ± SEM. Two-way ANOVA, *p<0.05, **p<0.01

The SAT is an additional useful test of anxiety-like behavior because unlike the EPM, it does not have a neutral junction region. Nevertheless, there were also no genotype effects, interaction effects, or genotypic differences in the stratified analyses of any behaviors measured in the SAT in either sex (**Figure 6D-G**). There were significant sex effects for average velocity (F_1,44_=7.59, p=0.0085), total distance traveled (F_1,44_=7.37, p=0.0094), white segment entry frequency (F_1,44_=4.49, p=0.0398), duration (F_1,44_=6.44, p=0.0148), and latency (F_1,44_=7.11, p=0.0107) as well as dark gray segment duration (F_1,44_=7.13, p=0.0106) (**Figure 6D-G**). Female rats traveled faster and farther than male rats, and they entered the white segment more often and more quickly while spending less time in the dark gray segment (**Figure 6D-G**).

### 4.7 Adcy3^mut/mut^ males and females both have altered feeding behavior in the NSF

In the NSF, due to different experimental conditions between sexes (see Methods 3.4.7), we stratified the data by sex and conducted analyses separately for males and females. Adcy3^mut/mut^ females spent significantly less time eating than WT females (t_19_=2.44, p=0.0249), indicating increased anxiety-like behavior (**Figure 7A**). Adcy3^mut/mut^ females also had a significantly higher latency to eat than WT females (t_18_=2.73, p=0.0136) (**Figure 7C**).

**Figure 7.**
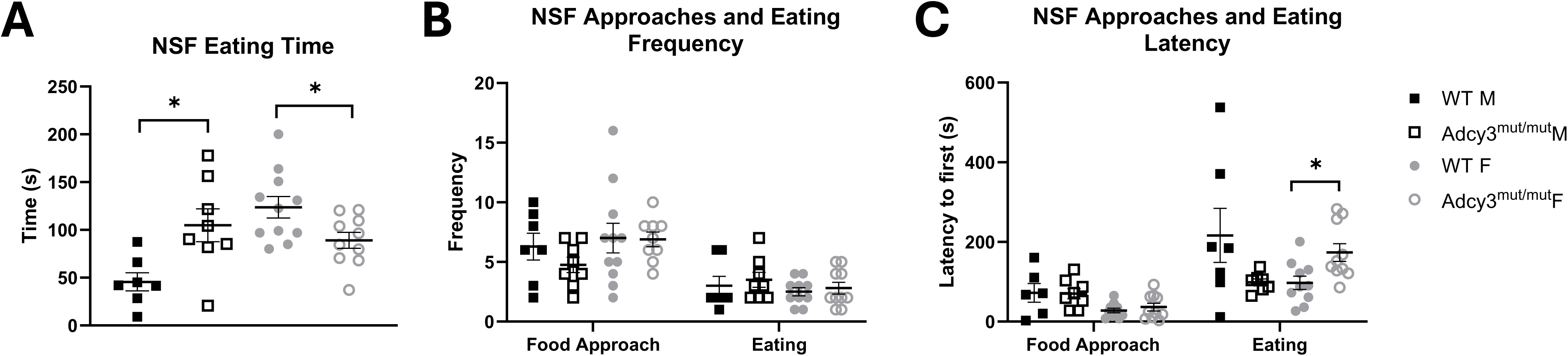
Adcy3^mut/mut^ behavior in the novelty-suppressed feeding test (NSF). **(A)** Adcy3^mut/mut^ males (M) spend significantly more time eating than wild-type (WT) M in the NSF, while Adcy3^mut/mut^ females (F) spend significantly less time eating than WT F. F also spend significantly more time eating than M. **(B)** No significant differences between Adcy3^mut/mut^ and WT rats in food approach frequency or eating frequency. **(C)** F have a significantly lower food approach latency than M, but Adcy3^mut/mut^ F have increased eating latency relative to WT F. N=7-11 due to removal of rats that did not eat at all during the test. Mean ± SEM. T-test or two-way ANOVA, *p<0.05, **p<0.01

Interestingly, Adcy3^mut/mut^ males spent significantly *more* time eating than WT males (t_13_=2.90, p=0.0125) (**Figure 7A**). Many of the Adcy3^mut/mut^ males had a shorter latency to eat than the WT males, although this was not significant, likely due to the high variation in WT males (**Figure 7C**). There were no significant differences between genotypes in the stratified analyses in eating frequency, food approach frequency, or food approach latency (**Figure 7B-C**).

### 4.8. Adcy3^mut/mut^ males, but not females, have increased serum leptin levels

There was a significant effect of genotype for serum leptin (F_1,39_=8.25, p=0.0066) with no sex or interaction effects. When separated by sex, Adcy3^mut/mut^ males had significantly higher serum leptin than WT males (t_17_=3.83, p=0.0013) with no significant differences in females (**Figure 8**).

**Figure 8.**
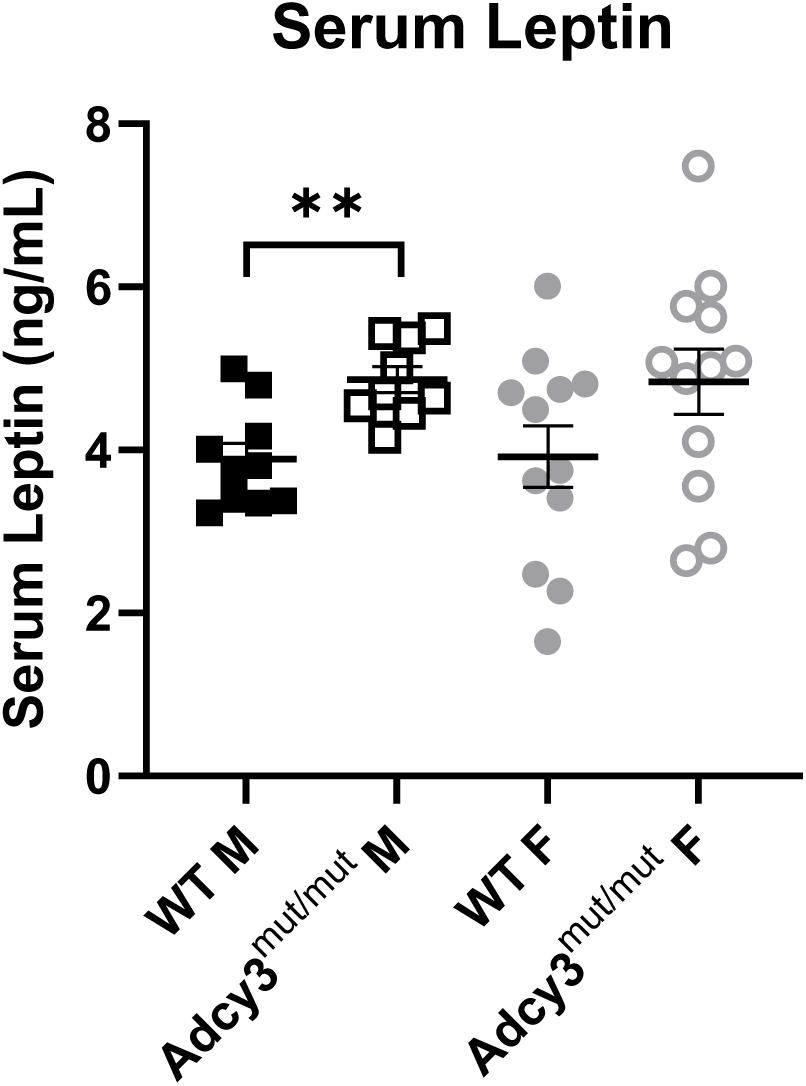
Serum leptin in Adcy3^mut/mut^. Adcy3^mut/mut^ males (M) have significantly higher serum leptin levels than wild-type (WT) M. There are no significant differences in serum leptin between Adcy3^mut/mut^ and WT females (F). Mean ± SEM. T-test, **p<0.01

### 4.9 Adcy3^mut/mut^ rats have decreased cAMP production in the hypothalamus with no differences in membrane or ciliary localization of ADCY3 protein

We assessed cAMP production in the hypothalamus to determine if Adcy3^mut/mut^ affects the enzymatic activity of ADCY3. Adcy3^mut/mut^ males had less cAMP in the hypothalamus than WT males, both with (t_14_=1.82, p=0.0900) and without (t_12_=3.97, p=0.0019) forskolin stimulation (**Figure 9A**). Adcy3^mut/mut^ females also had significantly less cAMP in the hypothalamus than WT females with forskolin stimulation (t_13_=4.14, p=0.0012), but there were no genotypic differences in cAMP production in females without forskolin stimulation (**Figure 9A**).

**Figure 9.**
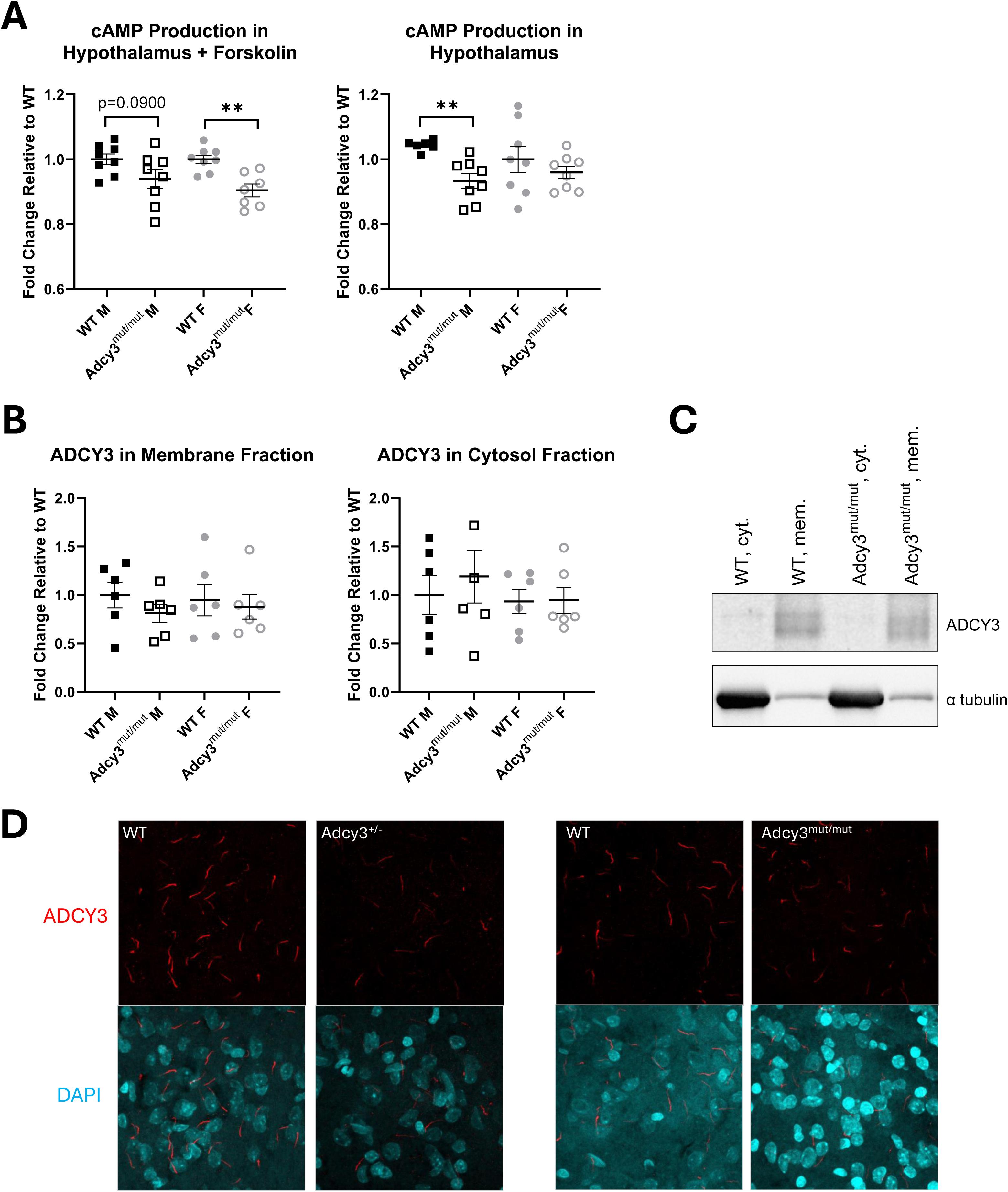
cAMP production and ADCY3 membrane localization in Adcy3^mut/mut^. **(A)** Adcy3^mut/mut^ males (M) have less cAMP in the hypothalamus than wild-type (WT) M, both with and without forskolin stimulation. Adcy3^mut/mut^ females (F) have significantly less cAMP production than WT F only with forskolin stimulation. **(B)** No differences in membrane or cytosolic ADCY3 content between Adcy3^mut/mut^ rats and WT rats. **(C)** Representative Western blot image of ADCY3 in the membrane and cytosol fractions. **(D)** Representative images of ADCY3- and DAPI-stained sections of the arcuate nucleus of the hypothalamus in male rats. Adcy3^+/-^ rats show less ADCY3 staining qualitatively than WT rats, while Adcy3^mut/mut^ rats do not. Mean ± SEM. T-test, **p<0.01

We also assessed membrane and cytosolic ADCY3 content in the hypothalamus to determine if Adcy3^mut/mut^ alters plasma membrane localization of the ADCY3 protein. There were no significant differences in membrane or cytosolic fraction ADCY3 content in Adcy3^mut/mut^ males or females relative to WT rats (**Figure 9B-C and Figure S2**). There were also no significant differences in membrane or cytosolic fraction ADCY3 content between genotypes in rats on a HFD (**Figure S3**).

Finally, we used immunofluorescence to qualitatively assess localization of ADCY3 to the neuronal cilia in WT and Adcy3^mut/mut^ male rats because ADCY3 has previously been identified as a critical protein for ciliary signaling.^31^ We used a genetic knockout model that we have previously reported has ∼50% expression of *Adcy3* (Adcy3^+/-^) for comparison.^37^ In the arcuate nucleus of the hypothalamus, Adcy3^+/-^ males showed less ADCY3 staining than WT males, but this difference was not present in Adcy3^mut/mut^ males relative to WT males (**Figure 9D**). Similar results were observed in the paraventricular nucleus of the hypothalamus (data not shown). These results suggest that Adcy3^mut/mut^ does not dramatically alter the ciliary localization of the ADCY3 protein.

## 5 Discussion

In the current study, we confirm that Adcy3^mut/mut^ rats have increased fat mass compared to WT rats.^37^ We then expand upon the previous work, showing that Adcy3^mut/mut^ males and females have altered negative emotion-like behaviors compared to WT rats in severalbehavioral tests, with key sex differences. Specifically, Adcy3^mut/mut^ males show increased despair-like (FST) and anxiety-like (OFT) behaviors as well as what we interpret as increased food drive (sucrose splash test, NSF, and possibly SPT). Meanwhile, Adcy3^mut/mut^ females show only increased anxiety-like behaviors in the OFT and NSF, with no differences in the other tests. An increased food drive in the Adcy3^mut/mut^ males is also supported by increased serum leptin (a hormone involved in satiety signaling^45^ and emotional eating^46^) relative to WT males. Finally, Adcy3^mut/mut^ rats of both sexes have decreased cAMP production in the hypothalamus, demonstrating that the TM domain of ADCY3 plays a key role in its enzymatic function. These findings emphasize the important role of the TM domain of ADCY3 in regulating adiposity and behavior, as well as confirm the presence of sex differences in several emotion-like behaviors.

Importantly, we have replicated the adiposity phenotype that we previously reported in Adcy3^mut/mut^.^37^ Both Adcy3^mut/mut^ males and females had increased fat mass, although only Adcy3^mut/mut^ males had increased body weight in this study. Regardless, the increased fat mass in both sexes is consistent with previous studies showing that *Adcy3* knockout (KO) rats also have increased adiposity.^30–32^ Of note, in preliminary studies (not shown), we found that a HFD was required to observe adiposity in Adcy3^mut/mut^ females and that a HFD exacerbated increased adiposity in Adcy3^mut/mut^ males, hence our decision to use a HFD for these *in vivo* studies. There is existing literature showing that *Adcy3* KO mice, particularly females, require a HFD to display adiposity as well.^30^ While we do not currently know if a HFD is required to observe behavioral changes in Adcy3^mut/mut^, these findings highlight a potential *Adcy3*-by-diet interaction that may have relevance for human obesity and mental health. Future studies should examine if the same behavioral phenotypes are present when rats are on normal chow to determine if HFD is required for the behavioral abnormalities seen in Adcy3^mut/mut^ rats. Nevertheless, in the current study, these results validate that both adiposity and behavioral phenotypes can be caused by the same protein-coding mutation in this gene under HFD conditions.

While we observed altered behavior in both Adcy3^mut/mut^ males and females, we found sex differences in most behavioral tests. Similar to our previous work,^37^ Adcy3^mut/mut^ females showed increased anxiety-like behaviors in the OFT and the NSF compared to WT females. Adcy3^mut/mut^ females groom more in the OFT and have both a longer eating latency and less eating time than WT females in the NSF, behaviors which have all been previously interpreted as “anxiety-like”.^47^ Of note, the OFT results in the present study differ slightly from our previous work where we found that Adcy3^mut/mut^ females had significantly decreased center time and rearing relative to WT females.^37^ We expect that these differences may be due to differences in size and lighting conditions of the OFT apparatus. Additionally, we did not observe altered behavior in Adcy3^mut/mut^ females in many of the behavioral tests we conducted, including the splash test, SPT, FST, EPM, and SAT. Collectively, these findings indicate that this increase in anxiety-like behaviors in Adcy3^mut/mut^ females is mild and likely test-specific. Thus, while we did observe some significantly increased anxiety-like behaviors in Adcy3^mut/mut^ females in both studies, we conclude that this mutation’s impact on emotion-like behaviors in females is relatively minimal, especially compared to its effects in males.

Adcy3^mut/mut^ males also show altered emotion-like behaviors, with decreased vertical rearing in the OFT (anxiety-like^48^) and increased immobility in the FST (depression-like^42^). This pattern of increased emotion-like behaviors is consistent with what we have previously reported in Adcy3^mut/mut^ male rats.^37^ Increased depression-like and anxiety-like behaviors have also previously been reported in the literature in male mice that have decreased *Adcy3.*^33, 34^

However, Adcy3^mut/mut^ males also unexpectedly showed increased grooming in the sucrose splash test and increased eating in the NSF, two behaviors that are typically associated with decreased emotionality in rodents.^47, 49^ Importantly, we have previously shown that Adcy3^mut/mut^ male rats consume significantly more food than WT rats,^37^ partially explaining the increased adiposity. Both the sucrose splash test and the NSF involve a food or sucrose-based motivating component. Thus, we hypothesize that we have inadvertently captured the increased food drive of the Adcy3^mut/mut^ males in these two tests. This is a compelling finding, demonstrating that the increased food drive in Adcy3^mut/mut^ males may be strong enough to overcome any potential increases in despair/anxiety-like behaviors that are also caused by this *Adcy3* mutation. Given that the sucrose preference test also uses a sucrose motivating component, and that reduced interest in sucrose would typically indicate anhedonia (a facet of depression-like behavior^47^), it is possible that we have captured increased food drive in the sucrose preference test as well. Furthermore, it is also possible that we have captured a form of emotional eating^50^ or even increased reward-seeking in these tests. *Adcy3* is known to play a role in satiety signaling in the hypothalamus,^51^ but *Adcy3* is also highly expressed in the basal ganglia, a region that plays a role in reward-related signaling.^29^ *Adcy3*’s role in this region is currently unknown. Future studies will be necessary to elucidate the specific neuronal circuits driving this phenotype.

We did not observe any altered behavior in Adcy3^mut/mut^ males or females in the EPM or SAT. We speculate that this can be attributed to the qualties of the WKY background strain on which on we generated Adcy3^mut/mut^. WT WKY rats are more passive than other rat strains and show increased passivity in several behavioral tests, including decreased time in the open arms of the EPM relative to other rat strains.^42, 52^ Similarly, we show that the WKY rats (both WT and Adcy3^mut/mut^) spent most of their time in the junction of the EPM and the enclosed area of the SAT. Thus, we believe that a floor effect in WKY behavior is prohibiting us from identifying differences between WT and Adcy3^mut/mut^ for these two tests.

Finally, we investigated the molecular mechanisms driving these phenotypes with a focus on understanding the mechanisms driving altered behavior. Adcy3^mut/mut^ males, but not females, have increased serum leptin compared to WT rats. This finding, combined with the fact that Adcy3^mut/mut^ males also consume more food than WT males,^37^ suggests that Adcy3^mut/mut^ males are leptin resistant.^45^ We cannot determine from the present study if the leptin resistance is a cause or effect of the increased adiposity. Importantly, however, increased serum leptin has also been associated with increased emotional eating,^46^ potentially supporting our behavioral findings. Thus, increased leptin levels, when combined with increased food intake^37^ and what we interpret as emotional eating in the splash test and NSF, support the theory that we have captured increased food drive in the Adcy3^mut/mut^ males in the present study.

We also found that both Adcy3^mut/mut^ males and females have decreased cAMP production in the hypothalamus compared to WT rats with no changes in membrane or ciliary localization. This finding suggests that this *Adcy3* mutation decreases the enzymatic capacity of the ADCY3 protein, contributing to a decrease in the total hypothalamic cAMP pool and likely explaining the previously reported decreases in downstream signaling proteins (PKA, pAMPK and CREB).^37^ Here, we measured cAMP in samples from rats on a chow diet as a baseline measurement of how ADCY3 function is affected by this mutation. But in the future, it will also be important to assess cAMP production in HFD samples to determine if a HFD exacerbates this deficiency. ADCY3 is one of multiple ADCY isoforms expressed in the hypothalamus, all of which contribute to cAMP concentrations in that tissue.^29^ Therefore, this decrease in cAMP likely represents ADCY3’s portion of the total cAMP pool in this tissue, although it is also possible that the enzymatic capacity of ADCY3 is not completely ablated. Additionally, although Adcy3^mut/mut^ is not in the catalytic domain of the ADCY3 protein, the TM domains have been shown to play a key role in promoting interactions between the two catalytic domains to maintain enzymatic function.^38^ The present study demonstrates that TM domain mutations of ADCY3 can cause deficits in the catalytic activity of this protein, even without affecting membrane or ciliary localization.

## 6 Conclusions

These results confirm that Adcy3^mut/mut^ can influence both adiposity and emotion-like behaviors, supporting the idea that obesity and MDD may be linked genetically and through shared biological mechanisms.^28^ In addition to successfully replicating the phenotypes reported in our initial study of Adcy3^mut/mut^,^37^ we report a potentially novel food-seeking and/or emotional eating phenotype in Adcy3^mut/mut^ males in several behavioral tests, as well as leptin resistance. Furthermore, we demonstrate key sex differences in emotional behaviors in Adcy3^mut/mut^ rats, where this mutation influences behavior more strongly in Adcy3^mut/mut^ males than females. Finally, we report that Adcy3^mut/mut^ impairs the enzymatic activity of ADCY3 in both sexes, leading to decreased cAMP in the hypothalamus, which likely drives the observed phenotypes. Future studies will differentiate between homeostatic and hedonic drivers of food seeking in Adcy3^mut/mut^ males by identifying the neuronal circuits that drive this increased feeding behavior, and future studies will also investigate the causes of the observed sex differences. These findings confirm that *Adcy3* plays an important role in both adiposity and behavior and that the TM domain of ADCY3 is critical for enzymatic function, supporting the consideration of genetic factors in future studies of both mental health disorders and obesity.

## Acknowledgements

The authors would like to thank the Medical College of Wisconsin Rodent Model Resource and the Rat Models and Genotyping Service Center for their assistance.

This work is supported by grants to Leah C. Solberg Woods (R01 DK120667), Mackenzie K. Fitzpatrick (2T32 DA041349-06), Jeffrey L. Weiner (AA26117), and Rong Chen (R01 AA030676, R21 DA056857).

## Data Availability Statement

The data that support the findings of this study are available from the corresponding author upon reasonable request.

## References

1. Substance Abuse and Mental Health Services Administration. (2022). Key substance use and mental health indicators in the United States: Results from the 2021 National Survey on Drug Use and Health. Center for Behavioral Health Statistics and Quality, Substance Abuse and Mental Health Services Administration, HHS Publication No. PEP22-07-01-005, NSDUH Series H-57.

2. Liu Q, He H, Yang J, Feng X, Zhao F, and Lyu J. (2020). Changes in the global burden of depression from 1990 to 2017: Findings from the Global Burden of Disease study. J Psychiatr Res, 126, 134–140. doi:10.1016/j.jpsychires.2019.08.002

3. Goodwin RD, Weinberger AH, Kim JH, Wu M, and Galea S. (2020). Trends in anxiety among adults in the United States, 2008-2018: Rapid increases among young adults. J Psychiatr Res, 130, 441–446. doi:10.1016/j.jpsychires.2020.08.014

4. Marx W, Penninx B, Solmi M, et al. (2023). Major depressive disorder. Nat Rev Dis Primers, 9(1), 44. doi:10.1038/s41572-023-00454-1

5. Giacobbe P and Flint A. (2018). Diagnosis and Management of Anxiety Disorders. Continuum (Minneap Minn), 24(3, behavioral neurology and psychiatry), 893–919. doi:10.1212/con.0000000000000607

6. Norkeviciene A, Gocentiene R, Sestokaite A, et al. (2022). A Systematic Review of Candidate Genes for Major Depression. Medicina (Kaunas), 58(2). doi:10.3390/medicina58020285

7. Gottschalk MG and Domschke K. (2017). Genetics of generalized anxiety disorder and related traits. Dialogues Clin Neurosci, 19(2), 159–168. doi:10.31887/DCNS.2017.19.2/kdomschke

8. Luppino FS, de Wit LM, Bouvy PF, et al. (2010). Overweight, Obesity, and Depression: A Systematic Review and Meta-analysis of Longitudinal Studies. Archives of General Psychiatry, 67(3), 220–229. doi:10.1001/archgenpsychiatry.2010.2

9. Mannan M, Mamun A, Doi S, and Clavarino A. (2016). Prospective Associations between Depression and Obesity for Adolescent Males and Females-A Systematic Review and Meta-Analysis of Longitudinal Studies. PLoS One, 11(6), e0157240. doi:10.1371/journal.pone.0157240

10. Berk M, Köhler-Forsberg O, Turner M, et al. (2023). Comorbidity between major depressive disorder and physical diseases: a comprehensive review of epidemiology, mechanisms and management. World Psychiatry, 22(3), 366–387. doi:10.1002/wps.21110

11. Wium-Andersen MK, Wium-Andersen IK, Prescott EIB, Overvad K, Jørgensen MB, and Osler M. (2020). An attempt to explain the bidirectional association between ischaemic heart disease, stroke and depression: a cohort and meta-analytic approach. Br J Psychiatry, 217(2), 434–441. doi:10.1192/bjp.2019.130

12. Zhang F, Rao S, and Baranova A. (2022). Shared genetic liability between major depressive disorder and osteoarthritis. Bone Joint Res, 11(1), 12–22. doi:10.1302/2046-3758.111.Bjr-2021-0277.R1

13. Pan A, Keum N, Okereke OI, et al. (2012). Bidirectional association between depression and metabolic syndrome: a systematic review and meta-analysis of epidemiological studies. Diabetes Care, 35(5), 1171–80. doi:10.2337/dc11-2055

14. World Obesity Atlas. 2023; Available from: https://data.worldobesity.org/publications/?cat=19.

15. Pulit SL, Stoneman C, Morris AP, et al. (2019). Meta-analysis of genome-wide association studies for body fat distribution in 694 649 individuals of European ancestry. Hum Mol Genet, 28(1), 166–174. doi:10.1093/hmg/ddy327

16. Yengo L, Sidorenko J, Kemper KE, et al. (2018). Meta-analysis of genome-wide association studies for height and body mass index in ∼700000 individuals of European ancestry. Hum Mol Genet, 27(20), 3641–3649. doi:10.1093/hmg/ddy271

17. Pérez-Gutiérrez AM, Carmona R, Loucera C, et al. (2024). Mutational landscape of risk variants in comorbid depression and obesity: a next-generation sequencing approach. Mol Psychiatry, 29(11), 3553–3566. doi:10.1038/s41380-024-02609-2

18. Zhang H, Zheng R, Yu B, et al. (2024). Dissecting shared genetic architecture between depression and body mass index. BMC Med, 22(1), 455. doi:10.1186/s12916-024-03681-9

19. Tong T, Yu R, and Park T. (2019). α-Cedrene protects rodents from high-fat diet-induced adiposity via adenylyl cyclase 3. Int J Obes (Lond), 43(1), 202–216. doi:10.1038/s41366-018-0176-0

20. Hines LM and Tabakoff B. (2005). Platelet adenylyl cyclase activity: a biological marker for major depression and recent drug use. Biol Psychiatry, 58(12), 955–62. doi:10.1016/j.biopsych.2005.05.040

21. Fujita M, Richards EM, Niciu MJ, et al. (2017). cAMP signaling in brain is decreased in unmedicated depressed patients and increased by treatment with a selective serotonin reuptake inhibitor. Mol Psychiatry, 22(5), 754–759. doi:10.1038/mp.2016.171

22. Chen D, Wang J, Cao J, and Zhu G. (2024). cAMP-PKA signaling pathway and anxiety: Where do we go next? Cell Signal, 122, 111311. doi:10.1016/j.cellsig.2024.111311

23. Qiu L, LeBel RP, Storm DR, and Chen X. (2016). Type 3 adenylyl cyclase: a key enzyme mediating the cAMP signaling in neuronal cilia. Int J Physiol Pathophysiol Pharmacol, 8(3), 95–108.

24. Markham A. (2021). Setmelanotide: First Approval. Drugs, 81(3), 397–403. doi:10.1007/s40265-021-01470-9

25. Redei EE, Andrus BM, Kwasny MJ, et al. (2014). Blood transcriptomic biomarkers in adult primary care patients with major depressive disorder undergoing cognitive behavioral therapy. Transl Psychiatry, 4(9), e442. doi:10.1038/tp.2014.66

26. Redei EE, Ciolino JD, Wert SL, et al. (2021). Pilot validation of blood-based biomarkers during pregnancy and postpartum in women with prior or current depression. Transl Psychiatry, 11(1), 68. doi:10.1038/s41398-020-01188-4

27. Stergiakouli E, Gaillard R, Tavaré JM, et al. (2014). Genome-wide association study of height-adjusted BMI in childhood identifies functional variant in ADCY3. Obesity (Silver Spring), 22(10), 2252–9. doi:10.1002/oby.20840

28. Fitzpatrick M and Solberg Woods LC. (2024). Adenylate cyclase 3: a potential genetic link between obesity and major depressive disorder. Physiol Genomics, 56(1), 1–8. doi:10.1152/physiolgenomics.00056.2023

29. Devasani K and Yao Y. (2022). Expression and functions of adenylyl cyclases in the CNS. Fluids Barriers CNS, 19(1), 23. doi:10.1186/s12987-022-00322-2

30. Tong T, Shen Y, Lee HW, Yu R, and Park T. (2016). Adenylyl cyclase 3 haploinsufficiency confers susceptibility to diet-induced obesity and insulin resistance in mice. Sci Rep, 6, 34179. doi:10.1038/srep34179

31. Siljee JE, Wang Y, Bernard AA, et al. (2018). Subcellular localization of MC4R with ADCY3 at neuronal primary cilia underlies a common pathway for genetic predisposition to obesity. Nat Genet, 50(2), 180–185. doi:10.1038/s41588-017-0020-9

32. Wang Z, Li V, Chan GC, et al. (2009). Adult type 3 adenylyl cyclase-deficient mice are obese. PLoS One, 4(9), e6979. doi:10.1371/journal.pone.0006979

33. Chen X, Luo J, Leng Y, et al. (2016). Ablation of Type III Adenylyl Cyclase in Mice Causes Reduced Neuronal Activity, Altered Sleep Pattern, and Depression-like Phenotypes. Biol Psychiatry, 80(11), 836–848. doi:10.1016/j.biopsych.2015.12.012

34. Yang XY, Ma ZL, Storm DR, Cao H, and Zhang YQ. (2021). Selective ablation of type 3 adenylyl cyclase in somatostatin-positive interneurons produces anxiety- and depression-like behaviors in mice. World J Psychiatry, 11(2), 35–49. doi:10.5498/wjp.v11.i2.35

35. Pitman JL, Wheeler MC, Lloyd DJ, Walker JR, Glynne RJ, and Gekakis N. (2014). A gain-of-function mutation in adenylate cyclase 3 protects mice from diet-induced obesity. PLoS One, 9(10), e110226. doi:10.1371/journal.pone.0110226

36. Keele GR, Prokop JW, He H, et al. (2018). Genetic Fine-Mapping and Identification of Candidate Genes and Variants for Adiposity Traits in Outbred Rats. Obesity (Silver Spring), 26(1), 213–222. doi:10.1002/oby.22075

37. Fitzpatrick MK, Szalanczy A, Beeson A, et al. (2024). Protein-coding mutation in Adcy3 increases adiposity and alters emotional behaviors sex-dependently in rats. Obesity (Silver Spring). doi:10.1002/oby.24178

38. Gu C, Sorkin A, and Cooper DM. (2001). Persistent interactions between the two transmembrane clusters dictate the targeting and functional assembly of adenylyl cyclase. Curr Biol, 11(3), 185–90. doi:10.1016/s0960-9822(01)00044-6

39. Parent CA and Devreotes PN. (1995). Isolation of inactive and G protein-resistant adenylyl cyclase mutants using random mutagenesis. J Biol Chem, 270(39), 22693–6. doi:10.1074/jbc.270.39.22693

40. Planchez B, Surget A, and Belzung C. (2019). Animal models of major depression: drawbacks and challenges. J Neural Transm (Vienna), 126(11), 1383–1408. doi:10.1007/s00702-019-02084-y

41. Becker M, Pinhasov A, and Ornoy A. (2021). Animal Models of Depression: What Can They Teach Us about the Human Disease? Diagnostics (Basel), 11(1). doi:10.3390/diagnostics11010123

42. Redei EE, Udell ME, Solberg Woods LC, and Chen H. (2023). The Wistar Kyoto Rat: A Model of Depression Traits. Curr Neuropharmacol, 21(9), 1884–1905. doi:10.2174/1570159×21666221129120902

43. van Dam EA, van der Harst JE, ter Braak CJ, Tegelenbosch RA, Spruijt BM, and Noldus LP. (2013). An automated system for the recognition of various specific rat behaviours. J Neurosci Methods, 218(2), 214–24. doi:10.1016/j.jneumeth.2013.05.012

44. Deacon RM. (2013). The successive alleys test of anxiety in mice and rats. J Vis Exp(76). doi:10.3791/2705

45. Myers MG, Jr., Leibel RL, Seeley RJ, and Schwartz MW. (2010). Obesity and leptin resistance: distinguishing cause from effect. Trends Endocrinol Metab, 21(11), 643–51. doi:10.1016/j.tem.2010.08.002

46. Cassioli E, Rossi E, Squecco R, et al. (2020). Reward and psychopathological correlates of eating disorders: The explanatory role of leptin. Psychiatry Res, 290, 113071. doi:10.1016/j.psychres.2020.113071

47. Belovicova K, Bogi E, Csatlosova K, and Dubovicky M. (2017). Animal tests for anxiety-like and depression-like behavior in rats. Interdiscip Toxicol, 10(1), 40–43. doi:10.1515/intox-2017-0006

48. Knight P, Chellian R, Wilson R, Behnood-Rod A, Panunzio S, and Bruijnzeel AW. (2021). Sex differences in the elevated plus-maze test and large open field test in adult Wistar rats. Pharmacol Biochem Behav, 204, 173168. doi:10.1016/j.pbb.2021.173168

49. Deal AW, Seshie O, Lenzo A, Cooper N, Ozimek N, and Solberg Woods LC. (2020). High-fat diet negatively impacts both metabolic and behavioral health in outbred heterogeneous stock rats. Physiol Genomics, 52(9), 379–390. doi:10.1152/physiolgenomics.00018.2020

50. Konttinen H. (2020). Emotional eating and obesity in adults: the role of depression, sleep and genes. Proc Nutr Soc, 79(3), 283–289. doi:10.1017/s0029665120000166

51. Yeo GSH, Chao DHM, Siegert AM, et al. (2021). The melanocortin pathway and energy homeostasis: From discovery to obesity therapy. Mol Metab, 48, 101206. doi:10.1016/j.molmet.2021.101206

52. Pardon MC, Gould GG, Garcia A, et al. (2002). Stress reactivity of the brain noradrenergic system in three rat strains differing in their neuroendocrine and behavioral responses to stress: implications for susceptibility to stress-related neuropsychiatric disorders. Neuroscience, 115(1), 229–42. doi:10.1016/s0306-4522(02)00364-0

