## Supplement for "A mutation in the transmembrane domain of *Adenylate cyclase 3* impairs enzymatic function to cause sex-specific depression- and anxiety-like behaviors and food seeking in a rat model"

**Supplemental Figures**

**Figure S1. Non-adipose tissue weights at euthanasia in wild-type (WT) and Adcy3^mut/mut^.** There were no genotypic differences in the weights of any non-adipose tissues at necropsy (sac). Mean ± SEM. T-test

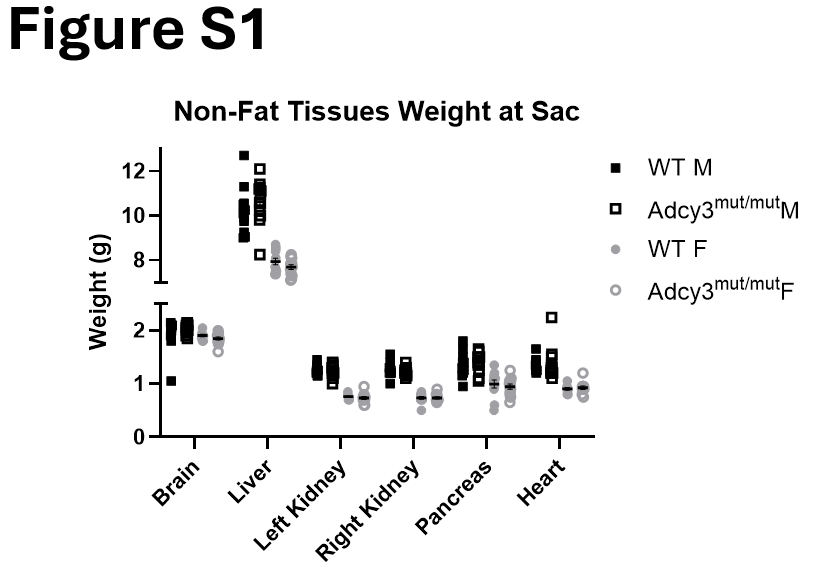

**Figure S2.** **Full western blot gels of ADCY3 in membrane and cytosolic fractions in the hypothalamus when animals were on chow diet, corresponding to representative Figure 9C.** Red boxes indicate the sections used for **Figure 9C**. Red lines on blot bands indicate saturation during imaging. Cyt: cytosolic fraction, mem: membrane fraction, WT: wild-type.

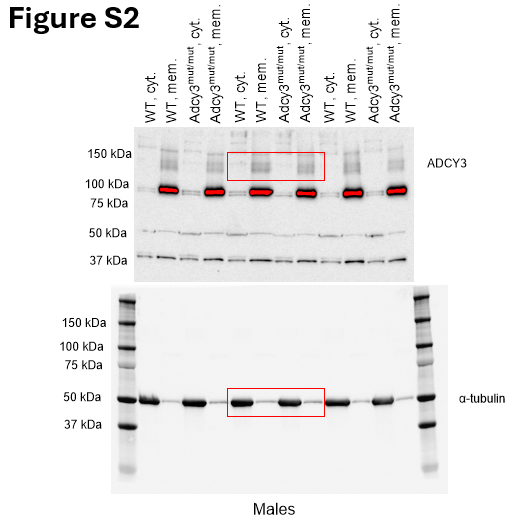

**Figure S3. ADCY3 membrane localization in Adcy3^mut/mut^ rats on a high-fat diet (HFD). (A)** No differences in ADCY3 content in the membrane or cytosolic fractions between Adcy3^mut/mut^ rats and wild-type (WT) rats on HFD. **(B)** Full Western blot gels of ADCY3 in membrane and cytosolic fractions in the hypothalamus when animals were on HFD. Red lines on blot bands indicate saturation during imaging. Cyt: cytosolic fraction, mem: membrane fraction. Mean ± SEM. T-test

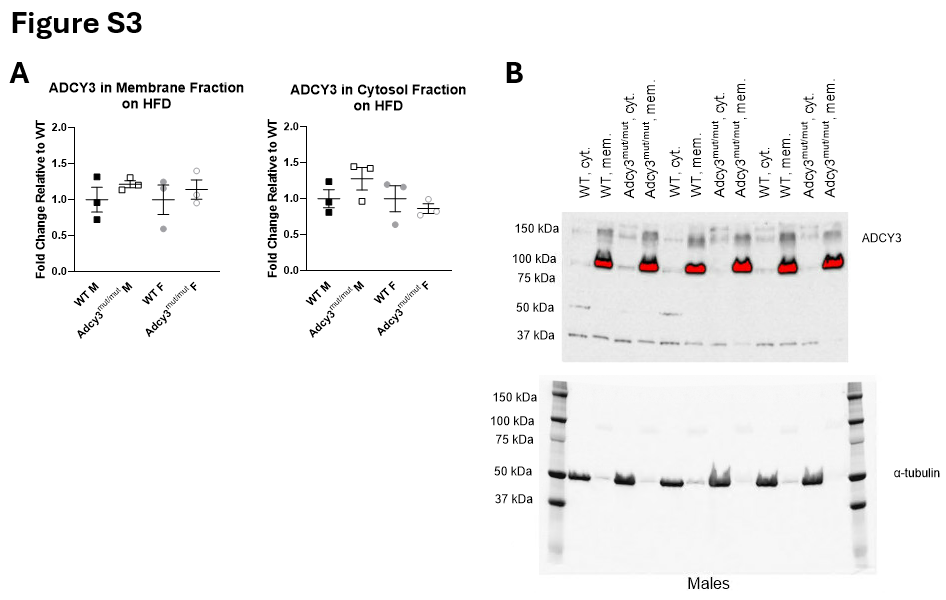
